# Development of a Stable Lyophilized Formulation of a Quadrivalent Frame-Shift Peptide Vaccine for Cancer Prevention in Lynch Syndrome

**DOI:** 10.1101/2025.08.14.670419

**Authors:** Shanker Gupta, Bijay Misra, Indu Javeri, Kaliappanadar Nellaiappan, Timothy Fouts, Robert H. Shoemaker

**Affiliations:** Chemopreventive Agent Development Research Group, Division of Cancer Prevention, NCI, NIH; CuriRx, Wilmington, MA; ABL, Inc. Rockville, MD

**Author notes:** Correspondence 9609 Medical Center Drive, Room 5E534, Rockville, MD 20850.

**Keywords:** cancer vaccine, neoantigen peptides, formulation development, lyophilization

## Abstract

A stable lyophilized formulation for a cancer vaccine containing quadrivalent frameshift neo antigen peptides TAFB(-1), AIM2(-1), HT001(-1) and TGFBR2(-1), was obtained through an iterative screening process. To develop a lyophilized formulation, the screening included evaluation of pH, stabilizers, bulking agents, and surfactants. The screening resulted in a scalable formulation and lyophilization process that co-formulated four peptides with 100 µg/ml each peptide in10 mM Histidine at pH 5.5, 260 mM trehalose, and 0.02% polysorbate 20 as a clinical formulation. Based on peptide content and peptide purity from a three-month stability study, the lyophilized formulation is estimated to be stable at ambient temperatures for 18 months.

## Introduction

The prophylactic vaccine for human papillomavirus, HPV infection represents a great achievement for cancer prevention. This vaccine has had a major impact in reducing cervical cancer^1^ as well as other cancer types. The vaccine targets highly immunogenic epitopes on the L1 capsid protein, presented on virus-like particles, and the current nonavalent vaccine against 9 different antigens protects about 90% of HPV-associated cancers. Development of vaccines for cancers not associated with viruses has been very challenging^1–2^. Some cancers are deficient in their ability to repair mismatched DNA which results in the accumulation of frameshift mutations in genes that contain microsatellite sequences, repeated sequences of DNA. Translation of these genes generates proteins with frameshift peptides induced by the mutations. These neoantigen peptides can provide a means for the immune system to target these microsatellite-unstable (MSI) cancers. Such frameshift peptides have been identified in four proteins, TAFB(-1), AIM2(-1), HT001(-1) and TGFBR2(-1) are being developed as vaccines to prevent/treat colorectal cancer, endometrial cancer, gastric cancer or small bowel cancer^23^ In the genomic era, it has become evident that individual patient’s tumors present unique mutational landscapes and associated neoantigens, creating problems for producing off-the-shelf vaccines. In the treatment setting, it has been possible to sequence tumors, identify mutations, predict immunogenicity of derived mutant peptides, synthesize them and thus generate personalized therapeutic vaccines. This sort of technological tour-de-force requires specialized facilities and a concerted effort for each patient vaccine ^3–19^.

The biology of Lynch Syndrome offers a different situation wherein frame-shift mutations recur in tumors from different patients. This enables an off-the-shelf vaccine composed of neoantigens derived from recurrent mutations. Investigators at the German Cancer Research Center in Heidelberg together with Micoryx have pioneered the development of multivalent frame-shift peptide vaccines and conducted an initial trial in patients with Lynch Syndrome and other microsatellite unstable colon cancers^6^. In this trial, individual peptides were combined with adjuvant (Montanide) and injected subcutaneously at multiple sites. The vaccine was shown to be safe and to generate an immune response.

To provide a precedent for use of such a vaccine to prevent cancer, an analogous multivalent peptide vaccine was created for use in a genetically engineered model of Lynch Syndrome. A quadrivalent frame-shift peptide vaccine adjuvanted with CPG was shown to prevent cancer in the mouse model, either alone or when combined with NSAIDs^2^. This vaccine was administered as a pool of four peptides subcutaneously.

To enable clinical testing of this concept, a quadrivalent frame-shift vaccine was designed based on truncated Micoryx peptides with the addition of a fourth peptide based on a very commonly recurrent (-1) frame-shift mutation in the gene for the TGFB receptor. Based on a series of studies in the mouse model, Hiltonol was selected as adjuvant and the intramuscular route was chosen for administration. Our goal was to provide a stable formulation in a single vial containing the four lyophilized frame-shift neoantigens for reconstitution with sterile water for injection and combination with adjuvant prior to injection. The overall objective is to manufacture a formulation of these quadrivalent frame-shift peptide antigens for use in Phase I clinical trials conducted by the National Cancer Institute (NCI). Asymptomatic carriers of DNA mis-match repair mutations (Lynch Syndrome carriers) will be the target population to be vaccinated and monitored for safety and immunogenicity. For clinical administration, the vialed and labeled quadrivalent peptide solution will be combined with adjuvant either at the bedside or at the hospital pharmacy. The four peptides are:

- TAFB(-1): NRGLKKKTILKKAGIGMCVKVSSIFFINKQKP
- AIM2(-1): VIKAKKKHREVKRTNSSQLV
- HT001(-1): RSNSKKKGRRNRIPAVLRTEGEPLHTPSVGMRETTGLGC
- TGFBR2(-1): KEKKSLVRLSSCVPVALMSAMTTSSSQKNITPAILT

The present paper describes the development of a vaccine formulation containing four tumor-specific neo-antigens for the intervention in MMR-deficient tumors in a Phase I clinical trial setting. A stable lyophilized formulation containing four peptides at 100 micrograms each of the four peptides that can be reconstituted and combined with an adjuvant either at the bedside or hospital pharmacy was developed.

The formulation development studies identified a stable buffer and pH for co-formulated peptides with negligible self-interaction. These studies identified stabilizers that are compatible with lyophilization process. Once the co-formulated fill solution was identified, lyophilization cycle development was performed, followed by a three-month stability study at 25°C and 40°C. These data are outlined below.

### Experimental Section

#### Materials

The peptides were manufactured for NCI at Polypeptide, Inc., CA. Histidine, trehalose, and polysorbate 20 were JT Baker Brand obtained from VWR Scientific, Radner, PA, Pfansthiel, Waukegan, IL, respectively. Water, HPLC and USP grade WFI was obtained from GE Lifesciences and PharmCo, AAPER, Connecticut.

## Methods

### Appearance

All samples were visually inspected for appearance using a 22W light source. The visual appearance was conducted to note the color, clarity, and presence of particulates.

### pH

The pH of peptide solution or placebo was measured using a Thermo Scientific, Orion Star Model A 211 pH meter equipped with a Ross PerpHecT microelectrode, Model 8220BNWP.

### Osmolality

Osmolality measurements were performed using a Model 3320 freezing-point micro-osmometer, equipped with a 20 µL Ease Eject™ Sampler.

### Reverse-phase chromatography

The Reverse-phase high-performance liquid chromatography (RP-HPLC) method was used to separate all 4 peptides and the corresponding impurities. The YMC Triart C18; 150×2mm, 1.9µm, 120Å PN: TA12SP9-1502PT was used with 0.1 % TFA in water and 0.1% TFA in acetonitrile mobile phases with a gradient elution for 45 minutes. The flow rate was maintained at 0.26 mL/min. The RP-HPLC method is used to determine peptide purity, impurities and peptide content. The peptide content is determined via an external standard method using the working reference standard.

### Flow Imaging Microscopy (FlowCam)

Subvisible particles in the sample were obtained with a FlowCam (Dynamic Fluid Imaging Systems, Inc.). The instrument was focused using 10 µm polystyrene beads at 3000 particles/mL NIST standard. An aliquot of 800 µL sample was aspirated at 0.08 mL/min, through a 100 µm x 1 mm flow cell and images of the particles were taken with a 10X optics system. The sample aliquot was imaged and particles per milliliter were calculated by operating Visual Spreadsheet software (Dynamic Fluid Imaging Systems, Inc.).

### Modulated Differential Scanning Calorimetry (MDSC)

MDSC analysis of the sample was performed using a Q2000 MDSC analyzer (TA instrument) to determine the glass transition temperature, Tg’, of the liquid formulation. An aliquot of liquid formulations, 30µL, was loaded into TZero pans (cat# T170103; TA systems) and crimped with TZero hermetic lids (T161025; TA systems). Empty TZero pan crimped with TZero hermetic lid was used as reference. The pans were ramped from 25°C to -60°C and from -60°C to 25°C at the ramp rate of 1°C/minute. The Tg’ was measured to set the primary drying temperature of lyophilization. Tg’ was obtained based on the thermal transition at the midpoint by the TA Universal Analysis software. A The primary drying temperature was set below the Tg’ at-30°C. As shown in Figure 8, lyophilization resulted a cake with no collapse.

### Simulated Shear Stress

To screen the effect of mechanical stress on liquid formulation, an aliquot of formulation was taken in Reacti-vial™ and stirred at 900 rpm for approximately 24 hours at 2-8°C. Unstable formulation develops haziness during this mechanical stress.

### Thermal Stress

Thermal stress was carried out at 25°C/60%RH and 40°C/75%RH in an incubator (Caron Model 60104 Environmental chamber) up to 14days for the screening studies. Both liquid and lyophilized formulations were subjected to thermal stress to identify the stable formulation. The lyophilized formulation obtained from the scale up batch was studied at 25°C/60%RH and 40°C/75%RH for three months.

### Lyophilization

Lyophilization was carried out using an FTS Lyo Star II lyophilizer. The formulation of peptides in, 10 mM Histidine, 260 mM trehalose, and 0.02% polysorbate 20, pH 5.5-6.0 was prepared, filtered through 0.2 µm filter, and filled 1.1 mL into 3 mL type 1 glass vial, partially stoppered with 13 mm FluroTec coated lyo stopper (West Pharmaceutical Services, Part # 19700034), The vials were placed in the lyophilizer, frozen at -50°C for 3 hours and dried at 75 millitorr pressure. Primary and secondary, drying were carried out at -30°C, and +20°C, respectively. The temperature ramp rate during the lyophilization cycle was maintained at 1°C/minute. The lyophilization cycle used for the scale up batch is listed in Table 1. The primary drying temperature was chosen based on MDSC determined Tg’

**Table 1:**
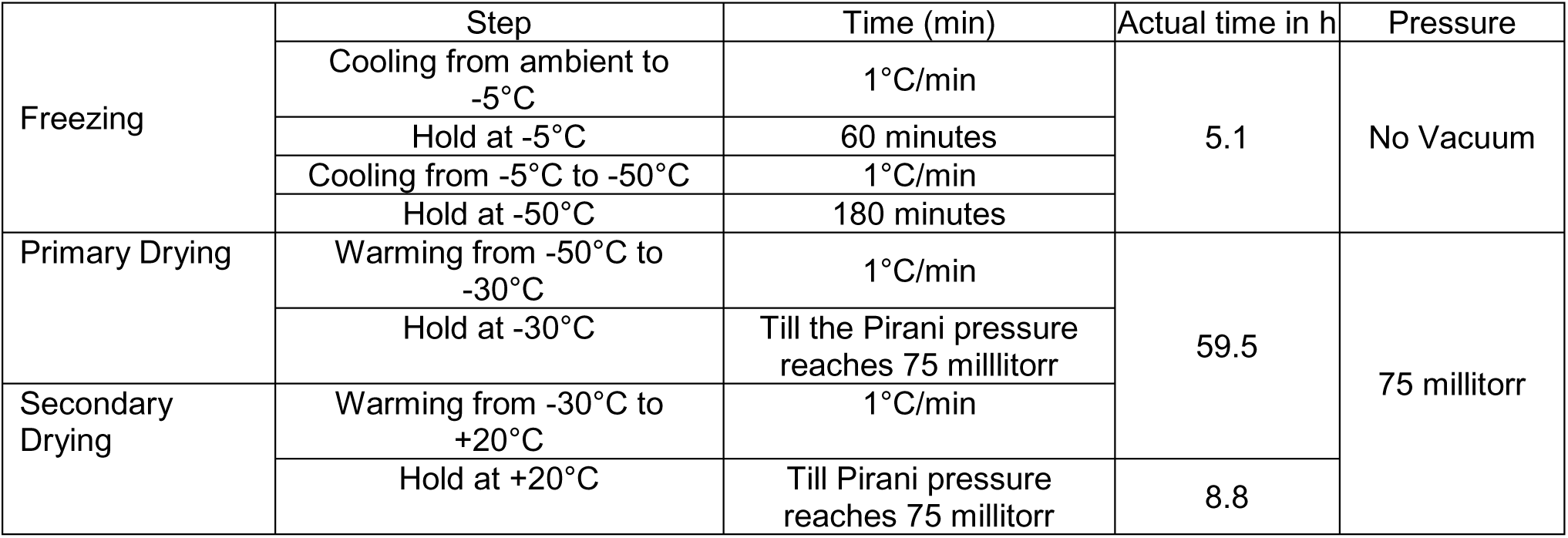
Lyophilization cycle development parameters for the lead formulation.

### Short-term stability study

Once an optimum formulation identified based on the stability of the peptides, a short-term stability study was conducted at 25°C/60%RH and 40 °C/75%RH for three months with sample pulls at 1-, 2- and 3-month periods. The formulation stability was evaluated by analyzing the quality attributes, such as the water content by KF, peptide purity and peptide content by reverse phase HPLC analysis. The peptide contents were determined via an external standard method.

### Moisture Content

The moisture content was determined using an 831 KF Coulometer with an 832 Thermoprep Drying Oven. The instrument was calibrated with a Hydranal water standard for KF-Oven (Sigma, 34784, Lot# SZBD 226AV). The vials with lyophilized cake were heated at 90°C in the drying oven and the water vapor generated was titrated coulometrically in Hydranal (Sigma, 34836, Lot# SZBE 2830V).

### Preparation of working reference standard

A 5 mL stock solution of each peptide was prepared at 0.4 mg/mL (based on net peptide content), in WFI and filtered through a 0.22 µm PES syringe filter. Equal volumes of each stock solution were combined to afford a working reference solution with100 µg/mL of each peptide. The working reference solution was aliquoted(100 µL) and stored at -80°C. The working reference standard was used to determine the peptide content in a sample by RP HPLC.

### Scale-up batch preparation

A scale-up batch of 1L of formulation was prepared.

### Reconstitution

The lyophilized cake was reconstituted with water for injection. The reconstitution volume was determined from the weight loss during the lyophilization. The water was added into the vial at center of the vial where the lyophilized cake dissolved immediately with no swirling. The reconstituted formulation was visually inspected for particulates. The subvisible particles were determined using Flow Imaging Microscopy. The peptide identification, purity, total impurities, and peptide content were determined by reverse phase-HPLC (Figure 1).

**Figure 1:**
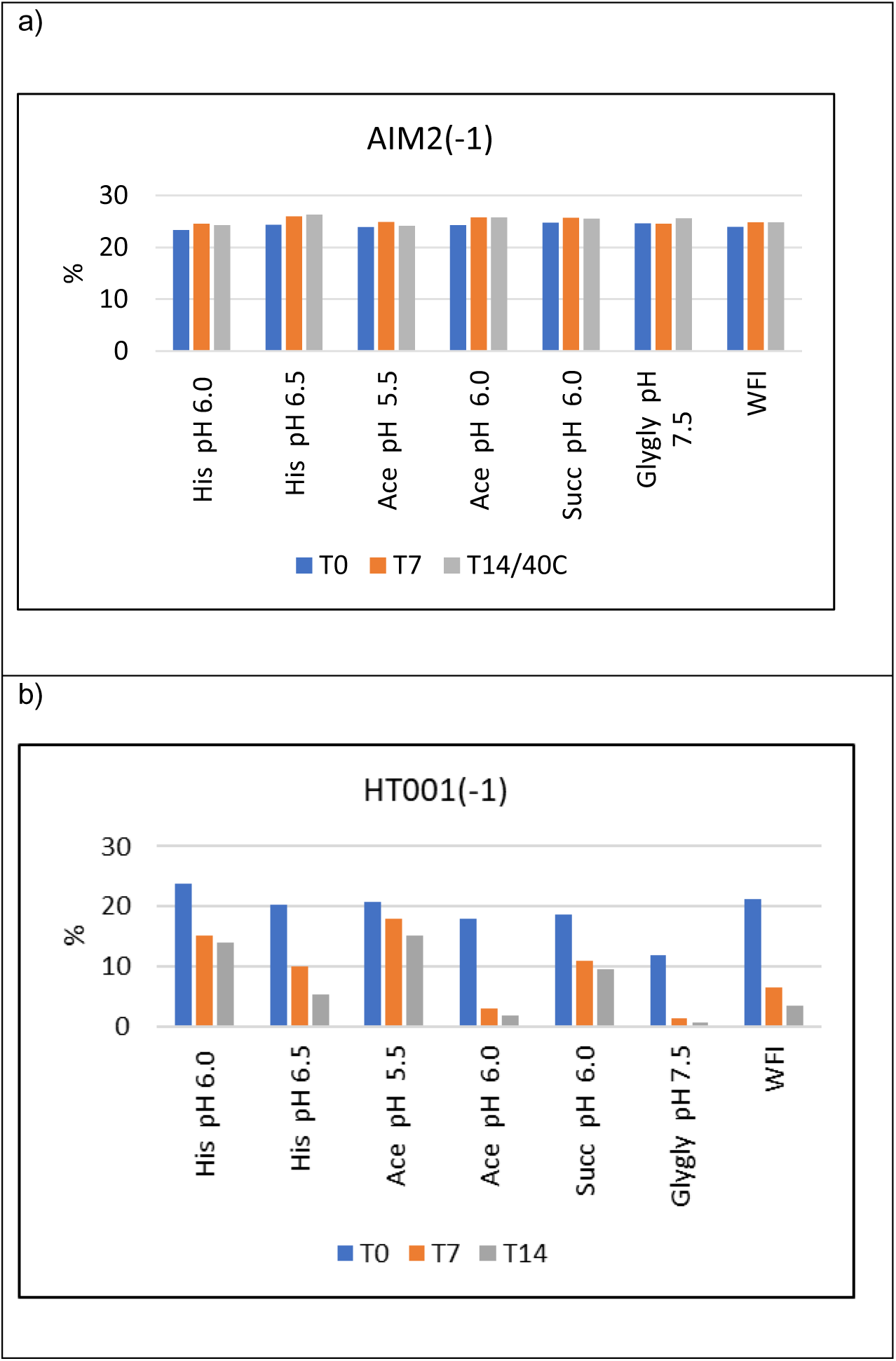
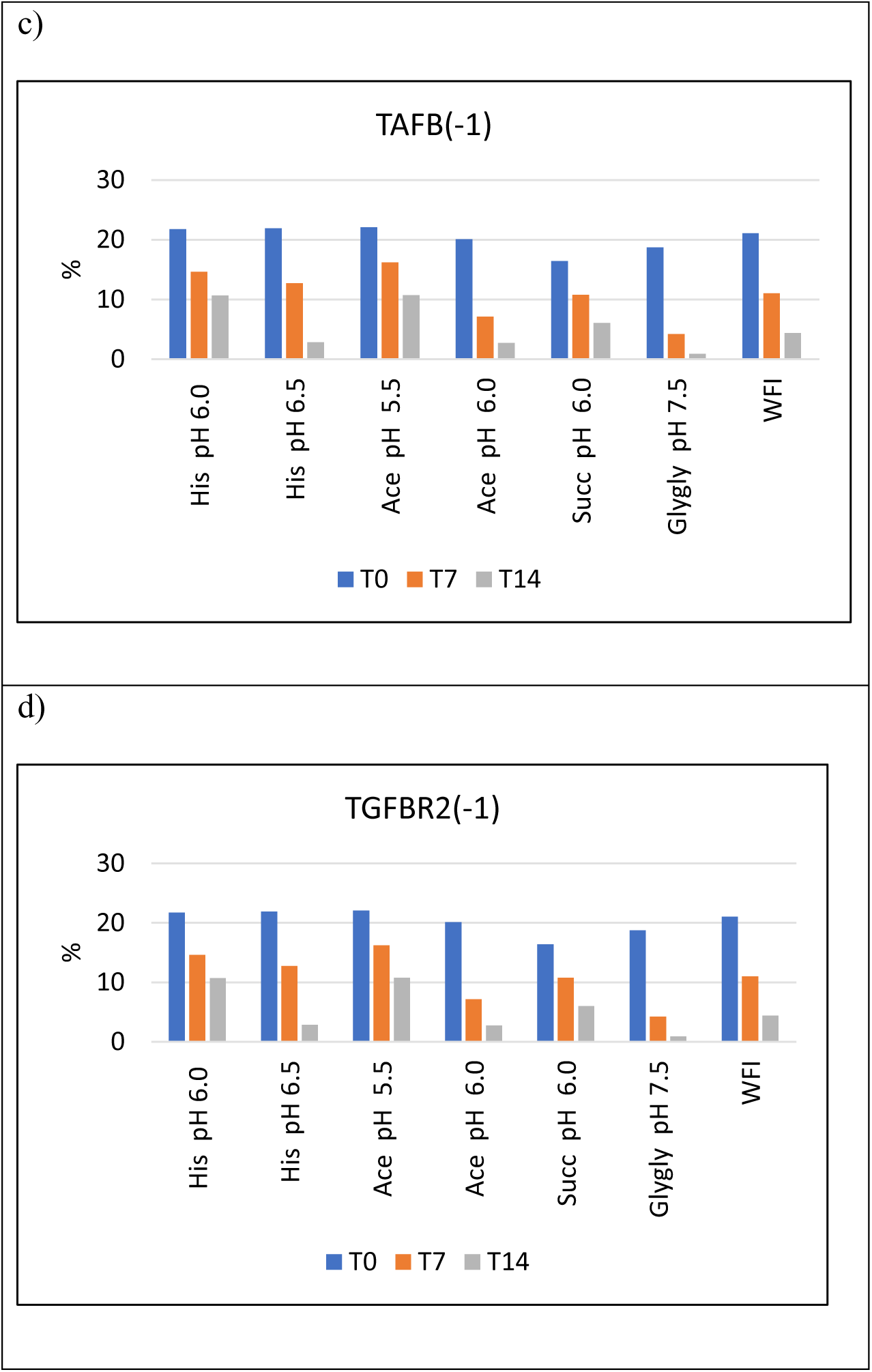
Buffer Screening – Peptide area percentage (n=3, CV=<5%) His – Histidine, Ace – Acetate, Succ – Succinate, Glygly - Glycinate

## Results

The formulation development for the quadrivalent peptide antigen solution was carried out in a systematic manner in four iterations at the target dose of 100 µg of each peptide. The screening process consisted of buffer and pH screening studies followed by stabilizers resulting in a fill solution that can be lyophilized. Freeze thaw and shear stress were performed for liquid formulations during the screening studies (data not shown). The screening of excipients was carried out simultaneously in liquid and lyophilized conditions. The selection of buffer-pH, excipient, stabilizer and surfactant were determined based on peptide stability at accelerated stability conditions, 40°C and 75% relative humidity over a two-week period. The lead formulation selected was carried forward for lyophilization cycle development. Since this product formulation is designed for Phase I clinical study, only targeted FDA approved excipients were considered, and the excipient bulking agent concentration is adjusted to be isotonic. The lyophilization primary drying temperature was determined using tg’ obtained by MDSC. A scale-up batch was produced to confirm the lyophilization parameters and subsequent stability of the lyophilized product.

### Different studies are outlined below

#### Buffer Evaluation Studies

The buffer evaluation studies consisted of a pH range 5.5 to 7.5 as shown in Figures 1 and 2. The buffer evaluation data for each peptide at 100 µg/mL indicated that except for AIM2(-1), the peptides are unstable in liquid formulations at 25°C and 40°C for 14 days (Figures 3). The total peptide impurities formed for the formulation in Histidine pH 6.0 and Acetate at pH 5.5 was observed to be the lowest. (Figures 3). Based on these data, formulation in histidine at pH 5.0-6.0 was selected for screening stabilizers as described below.

**Figure 2:**
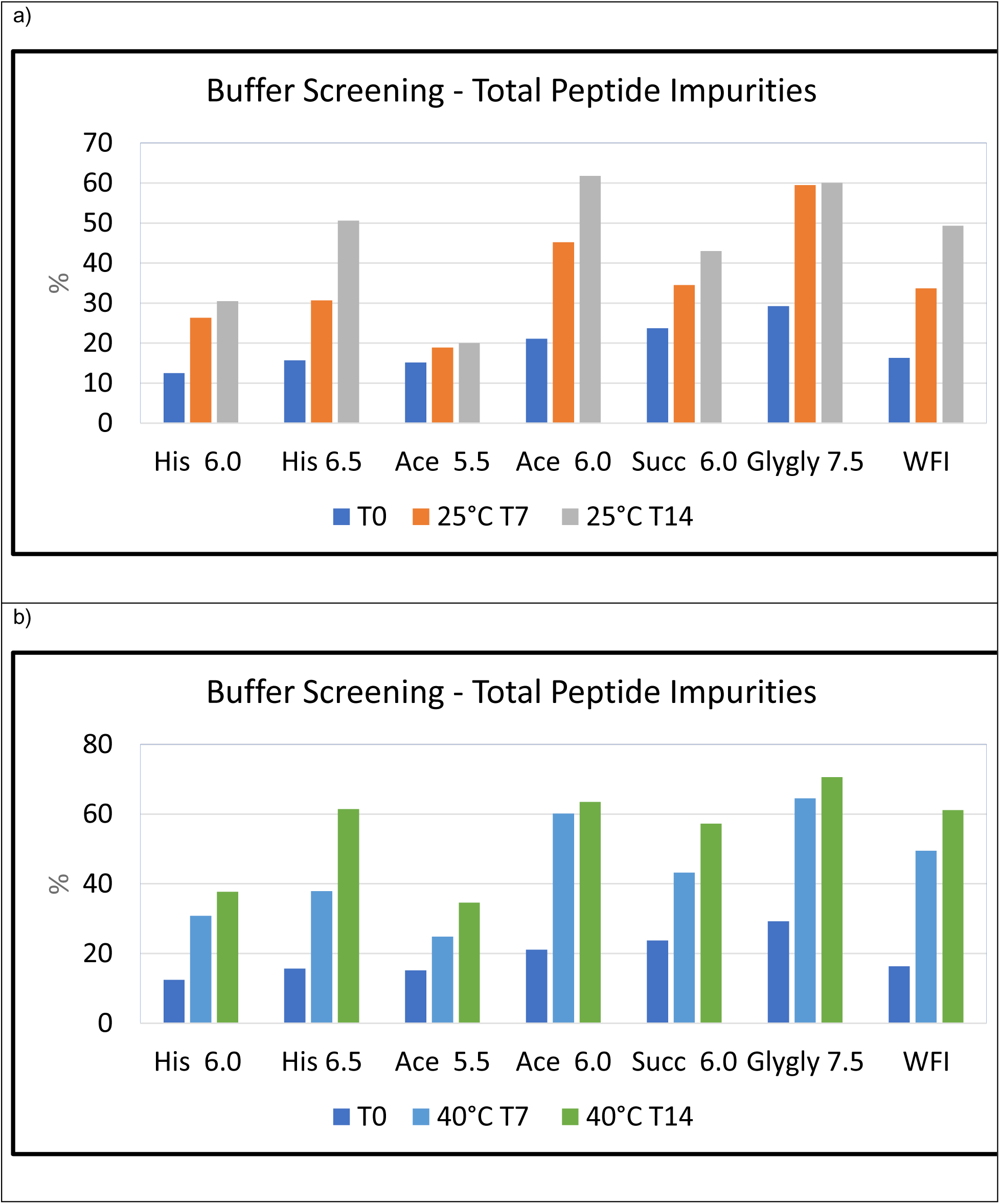
Buffer Screening – Total Peptide Impurities percentage (n=3, CV=<5%) His – Histidine, Ace – Acetate, Succ – Succinate, Glygly - Glycinate a)

**Figure 3:**
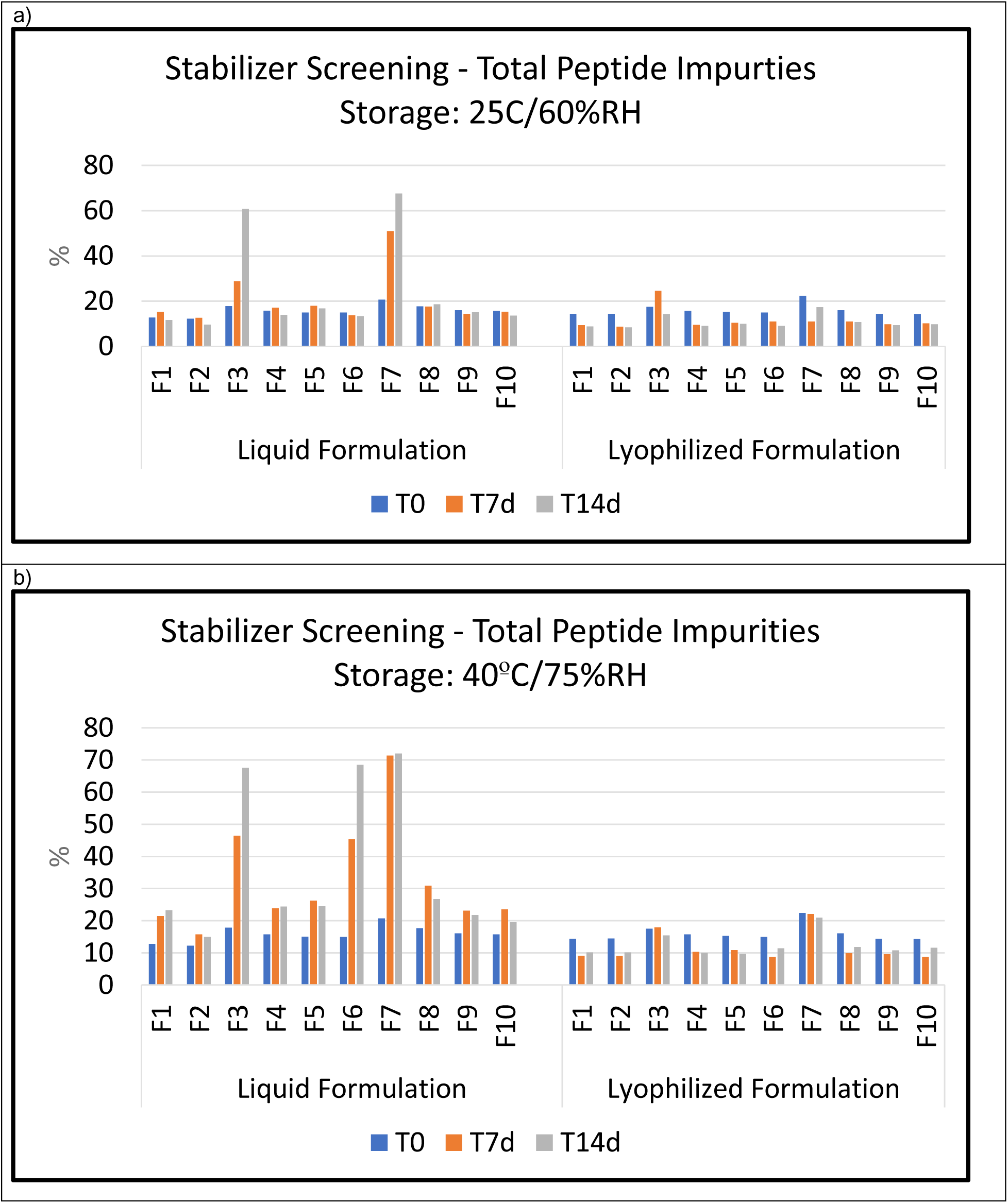
Stabilizer Screening – Total peptide impurities percentage (n=3, CV=<5%) , Suc – Sucrose, Tre – Trehalose, P188 – Poloxamer 188, PS20 and PS80 – Polysorbates 20 and 80, respectively

### Bulking Agent/Stabilizer Evaluation

The stabilizer screening was limited to targeted components with concentration suitable for a subcutaneous administration. Histidine buffers in the pH range 5.0-6.0 were evaluated in the presence of 260 mM trehalose or sucrose as stabilizers, bulking agents, or cryoprotectants and 0.1% poloxamer 188, 0.02% polysorbates 20 or 80 as surfactants. The goal was to develop a fill solution suitable for lyophilization and stable as a lyophilized product, so the screening was performed simultaneously as liquid and lyophilized formulations with accelerated thermal stress at 40°C. As shown in Figures 3, formulation in Histidine buffer at pH 5.0, with 260 mM Trehalose, and 0.02% polysorbate 20 exhibited relatively higher stability for all peptides with reduced formation of total peptide impurities in liquid formulations studied. The lyophilized formulations exhibited more stability compared to the corresponding liquid formulation. Among the lyophilized dosage formulations, the least impurities are seen in formulation containing 10 mM Histidine buffer at pH 5.0-6.0, 260 mM Trehalose and 0.02% polysorbates 20. The lyophilized formulation data in Table 3 also shows that polysorbates 80 destabilize the formulations. The formulations containing polysorbates 20 and poloxamer 188 are relatively more stable. However, the liquid formulation with P188 is relatively less stable compared to PS20 on thermal stress. The stability of the formulations containing trehalose and sucrose were comparable.

Based on the above data a formulation containing 10 mM histidine buffer at pH 5.0 – 6.0, 260 mM trehalose as a cryoprotectant, and 0.02% polysorbate 20 as a surfactant stabilizer was identified as the lead formulation for lyophilization feasibility study.

The lead formulation was characterized by its thermal properties using MDSC analysis (Figure 9) with the glass transition approximately at -30°C. Based on this data, the primary drying temperature for lyophilization was set at -30°C. The lyophilization parameters are listed in Table 1. The data indicate that the primary drying was completed in 60 hours, based on the Pirani pressure reaching the set chamber pressure of 75 millitorrs. The lyophilization cycle was completed in 74 hours with Pirani pressure at ≤75 millitorrs ensuring removal of the bound water. The formulation is subjected to thermal stress for four weeks and evaluated for peptide stability. The data shown in Figures 5-7 indicates that the co-formulated peptides in 10 mM Histidine at pH 5.5, 260 mM Trehalose, and 0.02% polysorbates 20 are stable with no significant increase in total peptide impurities or change in peptide area percentage by RP-HPLC in the lyophilized form under the stress conditions studied. The liquid formulation as fill solution was further evaluated for manufacturing conditions such as exposure to light, air, and agitation, and found to be stable for the production environment (data not shown).

**Figure 4:**
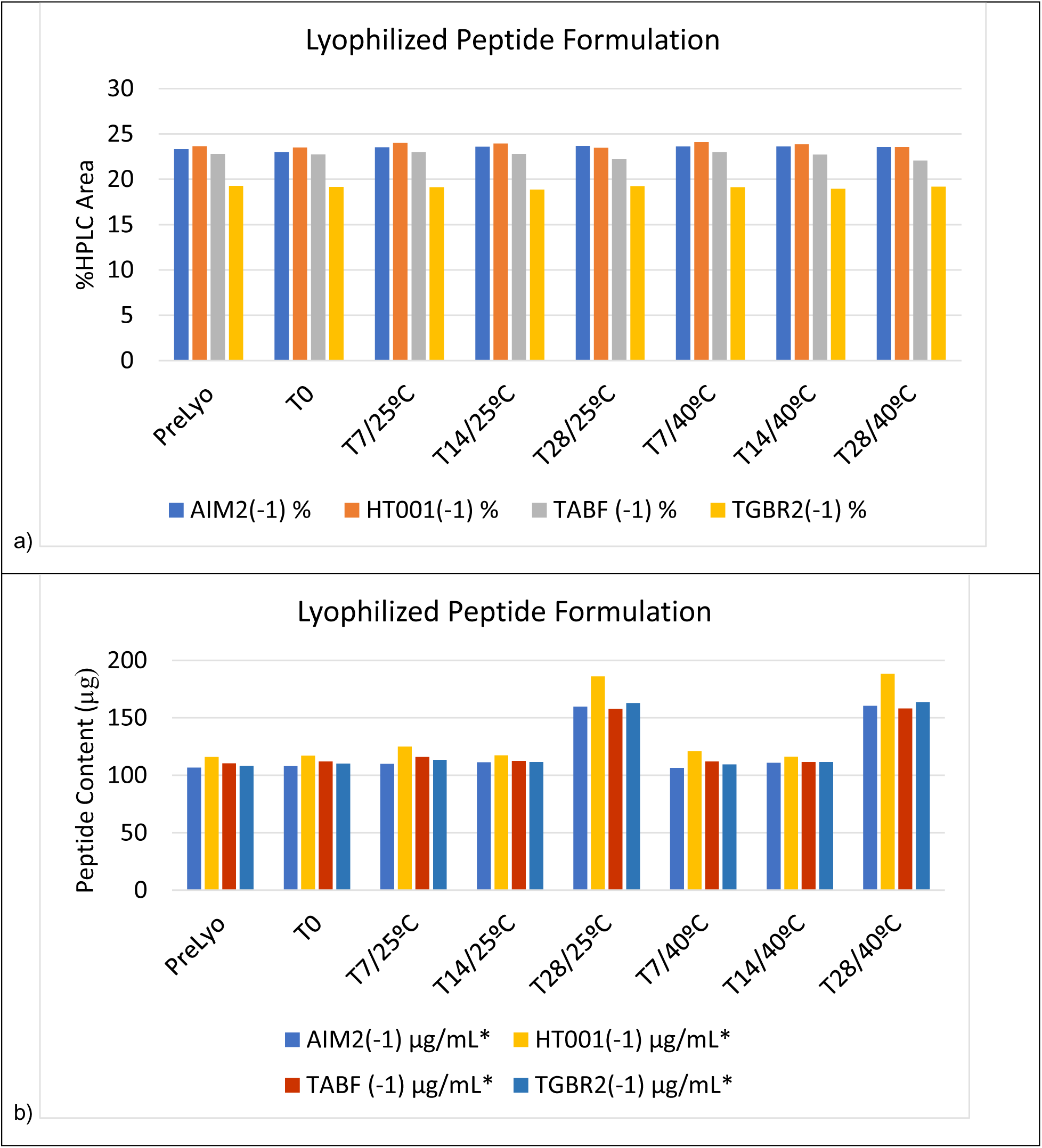
Lyophilization cycle development – Peptide area percent, content, and total impurities (n=3, CV=<5%) Suc – Sucrose, Tre – Trehalose, P188 – Poloxamer 188, PS20 and PS80 – Polysorbates 20 and 80, respectively The increased contents at T28 was due to reconstitution volume of 0.7 mL to accommodate the addition of adjuvant.

**Figure 5:**
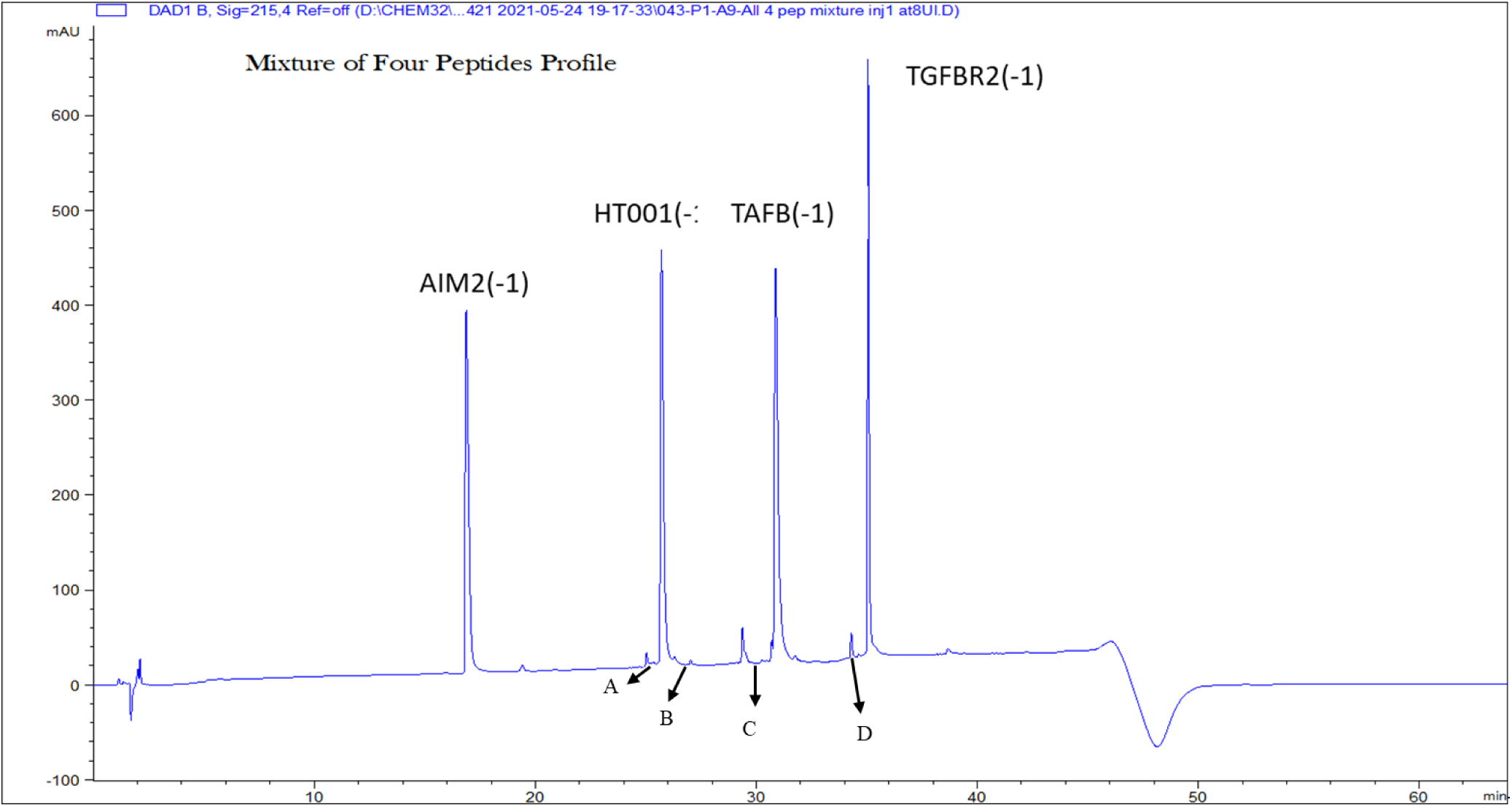
A Representative RP-HPLC Chromatogram of four peptides. The major peaks as shown were due to the four peptides and minor peaks were characterized as peptide fragments by LC-MS where fragments A and B belonged to HT001(-1), and fragments C and D belonged to TAFB (-1) and TGFBR2 (-1), respectively.

**Figure 6:**
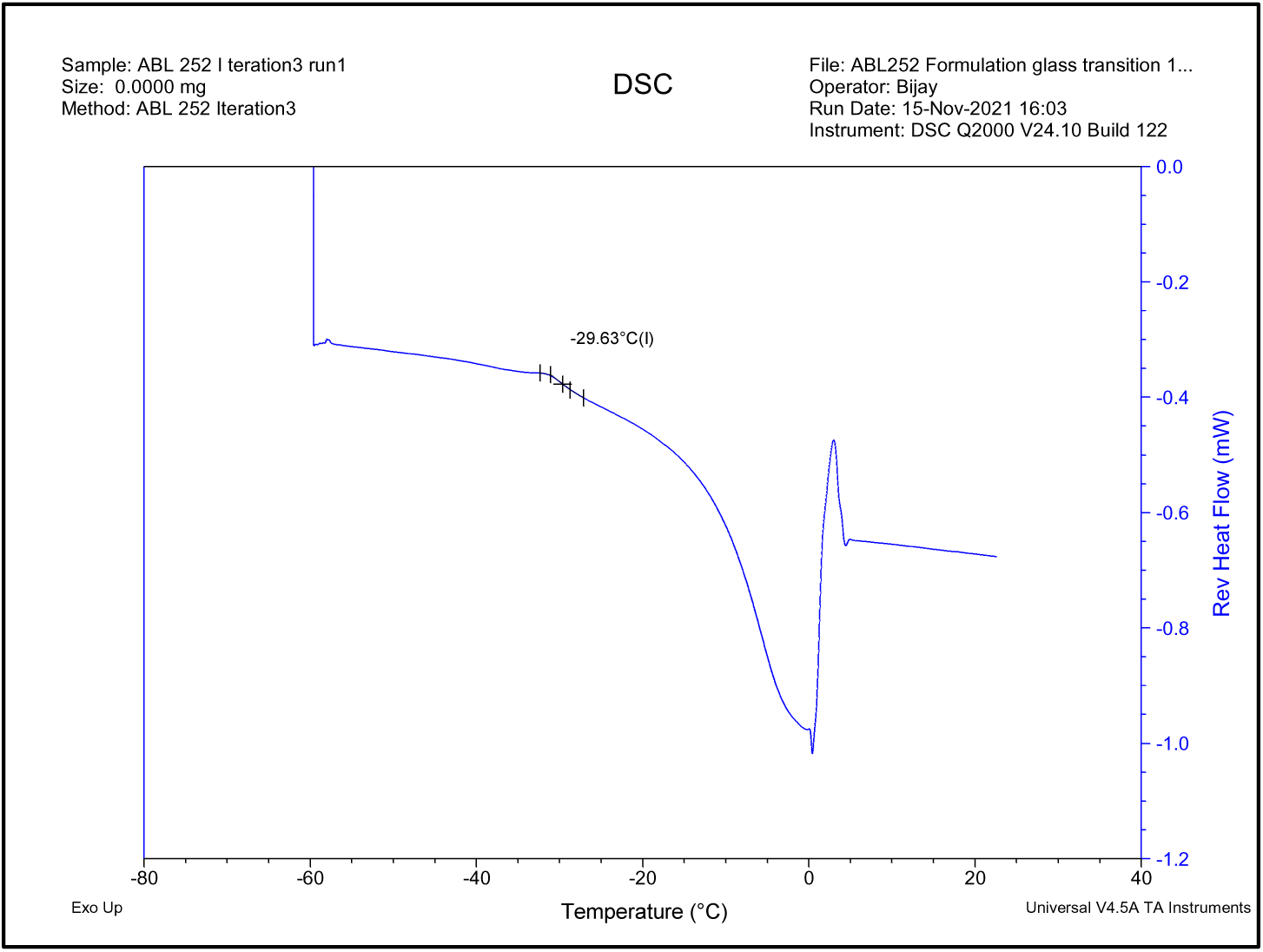
MDSC analysis of the lead liquid formulation – Tg’.

**Figure 7:**
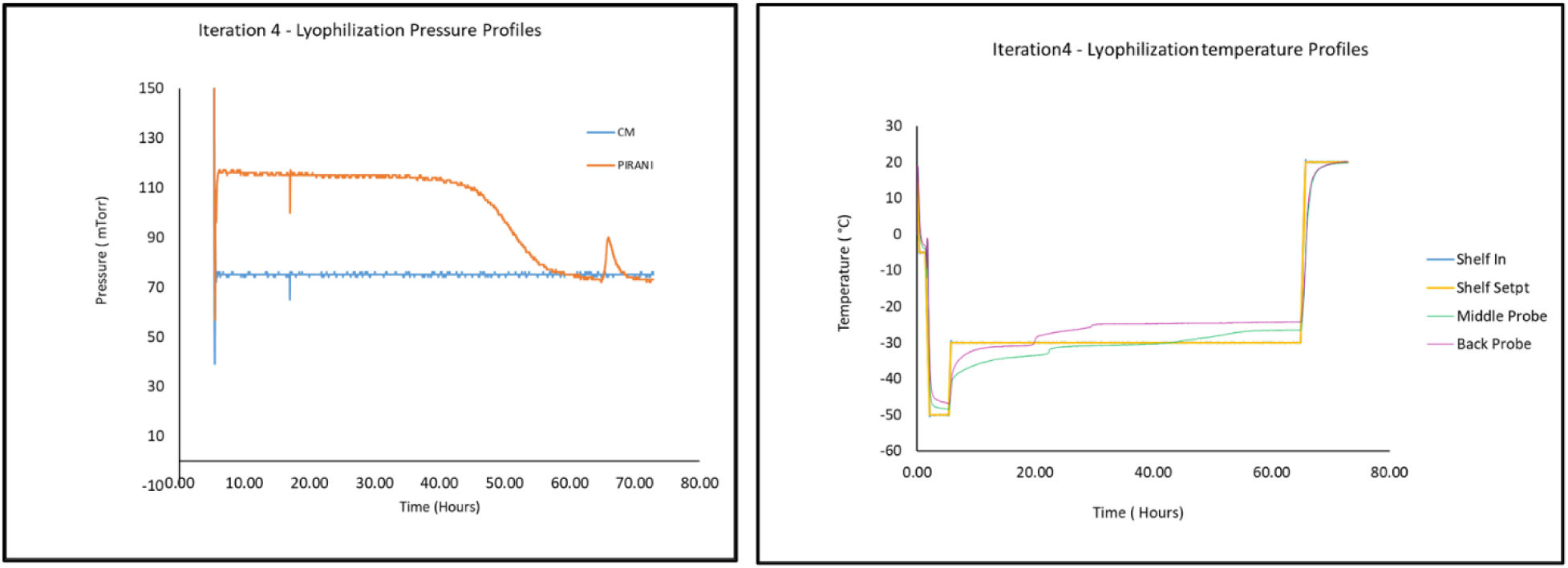
Lyophilization Feasibility - Pressure and Temperature profiles.

**Figure 8:**
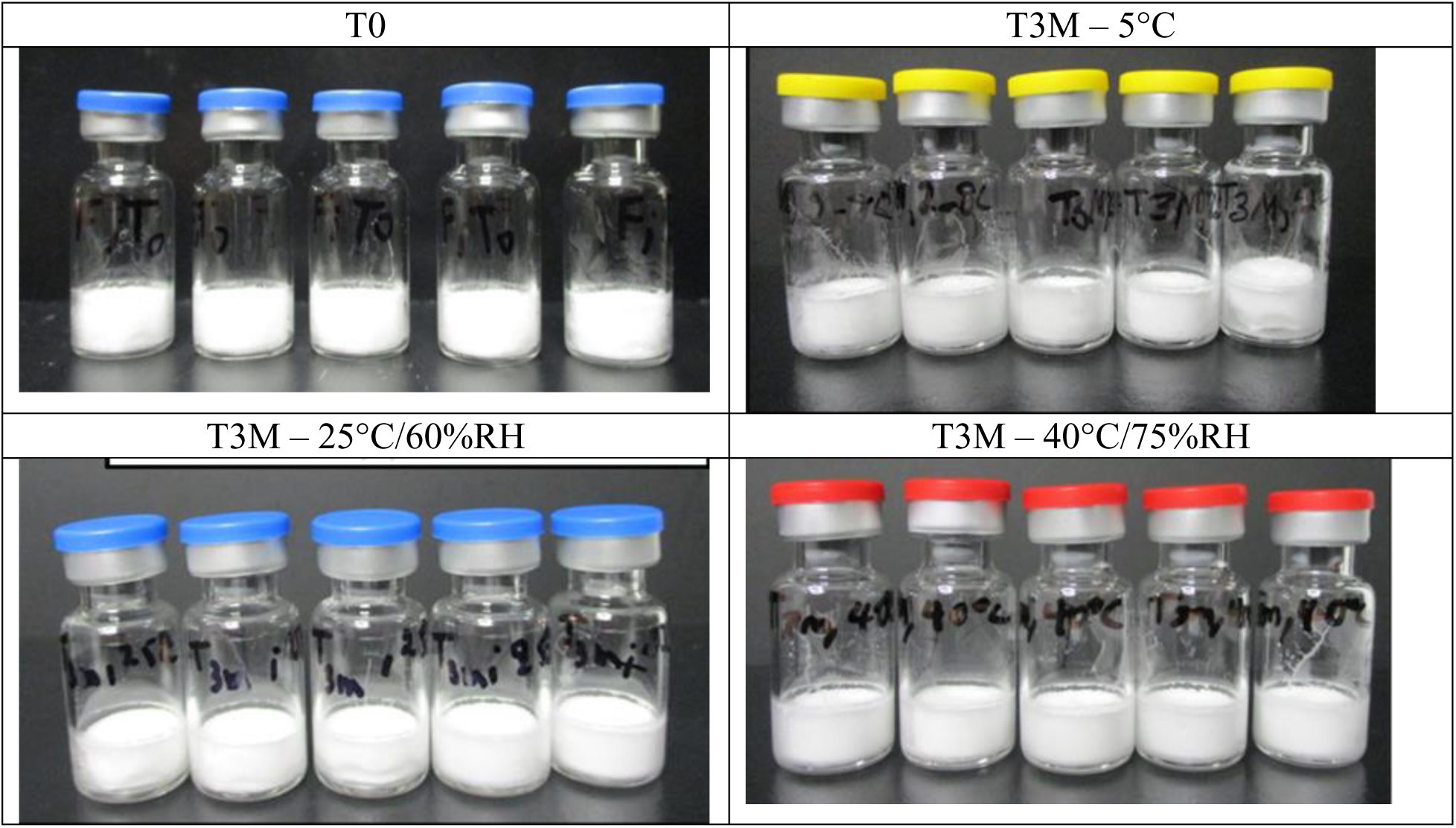
Three-month Stability Study - Appearance of Lyophilized peptide Formulation.

**Figure 9:**
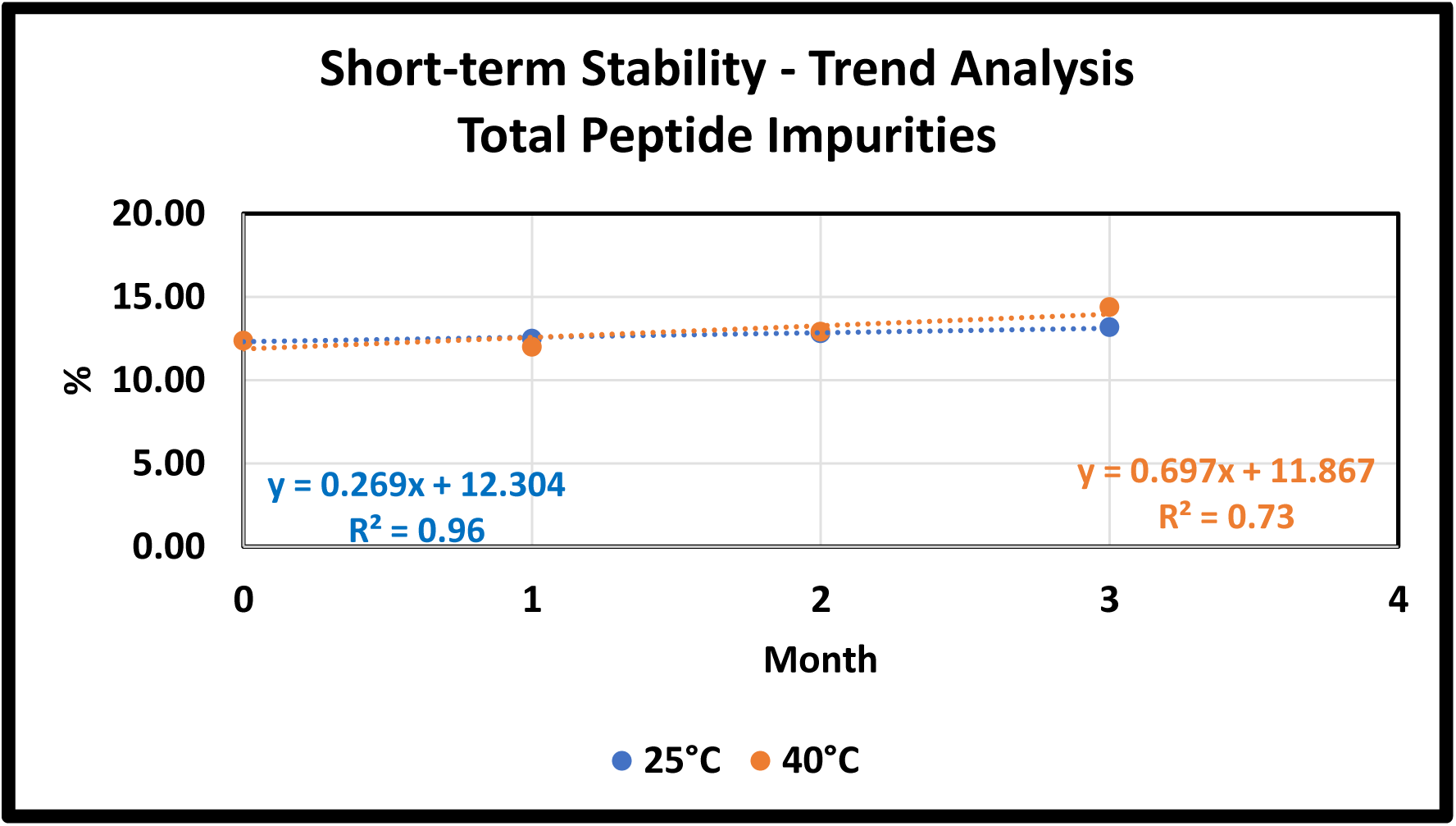
Effect of time and temperature on lyophilized co-formulated peptide: – Total Peptide impurity.

#### Lead Formulation – Manufacturability Evaluation

##### Preparation of a Scale-up Lyophilized Batch

A scale-up batch was prepared at a 1-liter scale as described in the material and method. This formulation was dispensed into vials and lyophilized. The lyophilization cycle completed in 120 hours (data not shown) and yielded a lyophilized product as white, intact, and without major collapse or shrinkage and comparable in appearance to the feasibility study.6.A 3-month stability study was carried out at 25°C/60%RH and 40°C/75%RH. The lyophilized cake appearance remained unchanged during the incubation at 25°C/60%RH and 40°C/75%RH for three months (Figure 11). The lyophilized cake was reconstituted in 1.02 mL water for injection resulting in a clear, colorless solution with no visible particles and 10+µm subvisible particles below 1000 p/mL. The reconstitution volume was determined from the loss in weight during the lyophilization.

The pH averaged 5.59± 0.05 (n=3) at 25°C/60%RH and 40°C/75%RH, indicating no change in pH values during the three-month study period. The osmolality averaged 308±3 mOsm/Kg, and the water content remained less than 2.5%.

Reverse phase analysis was carried out to evaluate peptide stability in the lyophilized formulation at 25°C/60%RH and 40°C/75%RH. All four peptide contents remained at 100 ± 20 µg/ml (data not shown). The peptide area percentage for AIM2(-1). HT001(-1), TAFB(-1) and TGFBR2(-1) were 22.8±0.4%, 22.4±0.2%, 21.7±0.2% and 19.3±0.2%, respectively. Although the peptide contents were approximately at the target of 100 µg/mL, the peptide area percent differed to each other due to the UV absorbance of the peptide due to the peptide sequence. The total impurities ranged from 11.99% to 14.39% with an average of 12.84±0.79% (n=7) and considered not significant. T28 samples reconstituted at 0.7 mL showed a higher osmolality at 490 mOsm/Kg due to the decrease of reconstitution volume from 1 mL into 0.7 mL. The corresponding slopes or increase in total impurities (Figure 12) were 0.3% and 0.7% percent per month, indicating the increase were about twice as fast at 40°C/75%RH compared to at 25°C/60%RH. To increase 5% impurity at 25°C/60%RH, at least 18 months would be required (Figure 12). If it is stored at 2-8°C, it is likely to be stable for more than 3 years (data not shown).

Overall, the lyophilization scale-up study resulted in a white intact lyophilized cake that was reconstituted in 30 seconds and the formulation quality attributes remained almost unchanged on storage at 25°C/60%RH and 40°C/75%RH for a period of three months. In conclusion, a stable lyophilized co-formulation of four frameshift peptides AIM2(-1), HT001(-1), TAFB(-1) and TGFBR2(-1) was achieved.

## Discussion

Ott et al (2017)^15^ demonstrated the feasibility, safety, and immunogenicity of a vaccine that targets up to 20 predicted personal tumor neoantigens. These antigens were formulated in DMSO, and sodium succinate at pH ≥ 5.5 so that the formulation is compatible with the adjuvant Hiltonol^®18^. These formulations were prepared in aqueous isotonic conditions and sterile filtered at ambient condition before sub-cutaneous administration to the patient ^13^.

In the current study, the four-frame shift peptide antigens aqueous formulation were demonstrated to be unstable and resulted degradation products. Therefore, the study goal was to develop a lyophilized formulation suitable for Phase I clinical trials that can be reconstituted at the bedside before its use.

A stable formulation with four peptides with a pI of approximately 9.0 was achieved by systematic evaluation of various candidate formulations. Peptide AIM2(-1) was found to be stable in all conditions studied, while the other three peptides were sensitive to pH and thermal conditions. The first step of the process included selecting a suitable buffer in combination with pH. The buffer at pH higher than 6.0 was found to be unsuitable with increased peptide degradation products. These were characterized by LC-MS and were found to be mostly peptide fragments. The pH 5.0 to 6.0 was observed to be more suitable with the formation of less peptide degradants. The excipients/stabilizers compatible with the peptides and for lyophilization were then evaluated. Trehalose and sucrose were used as bulking agents and cryoprotectants along with surfactants polysorbates 20, 80 and poloxamer 188. Polysorbates 80 was observed to destabilize peptides HT001(-1), TAFB(-1) and TGFBR2(-1).

All liquid formulations were unstable with increased peptide impurities at 40°C, and the major degradation pathway appears to be the hydrolysis of peptides in an aqueous solution. LC-MS analysis of the liquid formulations at accelerated storage conditions revealed that except peptide AIM2, all other peptides undergo hydrolytic degradation. AIM2 however, undergoes deamidation. There was no oxidation products observed for all four peptides (data not shown). The cryoprotectant trehalose was preferred over sucrose particularly in liquid formulation in pH 5.0 – 6.0 prevent potential Maillard reaction ^19^. The instability of formulation containing polysorbates 80 on thermal stress may be due to the oxidation of unstable oleic acid ester-containing double bonds, leading to the oxidation of peptides and formation of higher levels of impurities.

A three-month stability study of the lyophilized formulation at 25°C and 40°C confirms the compatibility of excipients with the four peptides studied. In this short-term stability study, the peptide content for each peptide remained at 100±20 µg/mL, the peptide area percentage remained within a standard deviation of 0.4% (n=7). The trend analysis shows that the formulation is stable at room temperature for more than 18 months based on an increase of impurity of ≤5%. Finally, a stable, ready-to-use lyophilized formulation containing four neoantigen peptides in a single vial was developed that can be readily reconstituted at the bedside prior to its administration

## Conclusion

The quadrivalent, frame-shift peptides AIM2(-1), HT001(-1) TGFBR2(-1), and TAF1B (-1), engineered in a novel lyophilized formulation with a 100-µg dose of each peptide, histidine (1.6 mg), trehalose dihydrate (98 mg), and polysorbates 20 (5 mg) is estimated to be stable for > 18 months @ 25°C. The lyophilized formulation is readily reconstituted in water, resulting in a particle-free clear solution with a pH of 5.5. This single-use lyophilized drug product stable at ambient conditions is suitable for the immunization of carriers of Lynch syndrome and other cancers exhibiting similar mutations.

## Acknowledgement

The authors thank Amina Soukrati for her assistance in the study.

## Supplementary Material

**Table 1:** Buffer Screening – Peptide area percentage (n=3, CV=<5%)

| Formulation ID* | AIM2(-1) |  |  | HT001(-1) |  |  | TAFB(-1) |  |  | TGFB2(-1) |  |  |
| --- | --- | --- | --- | --- | --- | --- | --- | --- | --- | --- | --- | --- |
|  | T0 | 40°C/75%RH |  | T0 | 40°C/75%RH |  | T0 | 40°C/75%RH |  | T0 | 40°C/75%RH |  |
|  |  | T7 | T14 |  | T7 | T14 |  | T7 | T14 |  | T7 | T14 |
| His pH 6.0 | 23.35 | 24.56 | 24.32 | 23.80 | 15.13 | 13.96 | 21.76 | 14.61 | 10.70 | 18.60 | 14.94 | 13.30 |
| His pH 6.5 | 24.38 | 26.01 | 26.29 | 20.34 | 10.14 | 5.44 | 21.91 | 12.72 | 2.85 | 17.69 | 13.19 | 4.00 |
| Ace pH 5.5 | 23.87 | 24.89 | 24.13 | 20.67 | 17.98 | 15.23 | 22.11 | 16.21 | 10.76 | 18.19 | 16.07 | 15.31 |
| Ace pH 6.0 | 24.27 | 25.77 | 25.76 | 17.88 | 3.05 | 1.94 | 20.13 | 7.14 | 2.72 | 16.62 | 3.92 | 6.10 |
| Succ pH 6.0 | 24.79 | 25.71 | 25.49 | 18.68 | 10.92 | 9.53 | 16.43 | 10.80 | 6.03 | 16.37 | 9.40 | 1.70 |
| Glygly pH 7.5 | 24.64 | 24.56 | 25.60 | 11.94 | 1.44 | 0.65 | 18.75 | 4.21 | 0.90 | 15.45 | 5.30 | 2.28 |
| WFI | 23.94 | 24.81 | 24.85 | 21.33 | 6.52 | 3.63 | 21.06 | 11.00 | 4.42 | 17.36 | 8.21 | 5.94 |

\*His – Histidine, Ace – Acetate, Succ – Succinate, Glygly - Glycinate

**Table 2:** Buffer Screening – Total Peptide Impurities percentage (n=3, CV=<5%)

| Formulation ID* | %Total Impurities |  |  |  |  |
| --- | --- | --- | --- | --- | --- |
|  | T0 | 25°C/60%RH |  | 40°C/75%RH |  |
|  |  | T7 | T14 | T7 | T14 |
| His pH 6.0 | 12.49 | 26.37 | 30.50 | 30.76 | 37.71 |
| His pH 6.5 | 15.68 | 30.67 | 50.64 | 37.94 | 61.42 |
| Ace pH 5.5 | 15.16 | 18.86 | 20.03 | 24.85 | 34.58 |
| Ace pH 6.0 | 21.10 | 45.18 | 61.78 | 60.12 | 63.49 |
| Succ pH 6.0 | 23.73 | 34.54 | 43.01 | 43.18 | 57.25 |
| Glygly pH 7.5 | 29.23 | 59.51 | 60.08 | 64.50 | 70.58 |
| WFI | 16.31 | 33.72 | 49.36 | 49.46 | 61.16 |

\*His – Histidine, Ace – Acetate, Succ – Succinate, Glygly - Glycinate

**Table 3:** Stabilizer Screening – Total peptide impurities percentage (n=3, CV=<5%)

| # | Formulation ID* | % Total Impurity (RP-HPLC) |  |  |  |  |
| --- | --- | --- | --- | --- | --- | --- |
|  |  | T0 | 25°C/60%RH |  | 40°C/75%RH |  |
|  |  |  | T7 | T14 | T7 | T14 |
| F1-Liq | His-Tre-Peptides-pH 6.0 | 12.79 | 15.19 | 11.74 | 21.46 | 23.29 |
| F2-Liq | His-Tre-Peptides-PS20-pH 5.0 | 12.23 | 12.63 | 9.70 | 15.74 | 14.97 |
| F3-Liq | His-Tre-Peptides-PS80-pH 6.0 | 17.83 | 28.74 | 60.75 | 46.44 | 67.52 |
| F4-Liq | His-Tre-Peptides-P188-pH 6.0 | 15.76 | 17.01 | 13.96 | 23.83 | 24.42 |
| F5-Liq | His-Suc-Peptides-pH 6.0 | 15.04 | 17.97 | 16.89 | 26.28 | 24.52 |
| F6-Liq | His-Suc-Peptides-PS20-pH 5.0 | 14.98 | 13.82 | 13.43 | 45.37 | 68.46 |
| F7-Liq | His-Suc-Peptides-PS80-pH 6.0 | 20.75 | 51.00 | 67.59 | 71.36 | 72.03 |
| F8-Liq | His-Suc-Peptides-P188-pH 6.0 | 17.72 | 17.61 | 18.62 | 30.87 | 26.75 |
| F9-Liq | His-Tre-Peptides-PS20-pH 5.5 | 16.05 | 14.41 | 15.08 | 23.14 | 21.80 |
| F10-Liq | His-Tre-Peptides-P188-pH 5.5 | 15.75 | 15.31 | 13.62 | 23.51 | 19.52 |
| F1-Lyo | His-Tre-Peptides-pH 6.0 | 14.43 | 9.45 | 8.85 | 9.10 | 10.16 |
| F2-Lyo | His-Tre-Peptides-PS20-pH 5.0 | 14.45 | 8.70 | 8.46 | 9.03 | 10.17 |
| F3-Lyo | His-Tre-Peptides-PS80-pH 6.0 | 17.51 | 24.60 | 14.23 | 17.89 | 15.45 |
| F4-Lyo | His-Tre-Peptides-P188-pH 6.0 | 15.74 | 9.58 | 9.12 | 10.32 | 9.99 |
| F5-Lyo | His-Suc-Peptides-pH 6.0 | 15.26 | 10.44 | 10.00 | 10.86 | 9.64 |
| F6-Lyo | His-Suc-Peptides-PS20-pH 5 | 15.00 | 11.04 | 9.12 | 8.80 | 11.42 |
| F7-Lyo | His-Suc-Peptides-PS80-pH 6.0 | 22.38 | 11.04 | 17.40 | 22.07 | 20.98 |
| F8-Lyo | His-Suc-Peptides-P188-pH 6.0 | 16.05 | 11.04 | 10.80 | 9.89 | 11.80 |
| F9-Lyo | His-Tre-Peptides-PS20-pH 5.5 | 14.43 | 9.80 | 9.45 | 9.60 | 10.81 |
| F10-Lyo | His-Tre-Peptides-P188-pH 5.5 | 14.30 | 10.22 | 9.80 | 8.82 | 11.60 |

\*Liq – Liquid formulation, Lyo – Lyophilized formulation, Suc – Sucrose, Tre – Trehalose, P188 – Poloxamer 188, PS20 and PS80 – Polysorbates 20 and 80, respectively.

**Table 4:** Lyophilization cycle development – Peptide area percent, content, and total impurities (n=3, CV=<5%)

| Attributes | PreLyo | 25C/60%RH |  |  |  | 40C/75%RH |  |  |  |
| --- | --- | --- | --- | --- | --- | --- | --- | --- | --- |
|  |  | T0 | T7 | T14 | T28 | T0 | T7 | T14 | T28 |
| AIM2(-1) % | 23.32 | 23.01 | 23.54 | 23.6 | 23.68 | 23.01 | 23.61 | 23.63 | 23.58 |
| HT001(-1) % | 23.65 | 23.52 | 24.04 | 23.96 | 23.47 | 23.52 | 24.1 | 23.85 | 23.57 |
| TABF (-1) % | 22.8 | 22.73 | 23.01 | 22.8 | 22.21 | 22.73 | 23.01 | 22.72 | 22.07 |
| TGBR2(-1) % | 19.28 | 19.16 | 19.14 | 18.87 | 19.25 | 19.16 | 19.12 | 18.94 | 19.20 |
| AIM2(-1) µg/mL* | 106.8 | 107.9 | 109.9 | 111.4 | 159.9 | 107.9 | 106.4 | 110.9 | 160.4 |
| HT001(-1) µg/mL* | 115.9 | 117.1 | 125.1 | 117.4 | 186.0 | 117.1 | 121.1 | 116.3 | 188.2 |
| TABF (-1) µg/mL* | 110.4 | 112.0 | 116.1 | 112.6 | 158.0 | 112.0 | 112.06 | 111.6 | 158.2 |
| TGBR2(-1) µg/mL* | 108.2 | 110.3 | 113.5 | 111.7 | 162.9 | 110.3 | 109.5 | 111.5 | 163.7 |
| Total Impurity % | 10.96 | 11.58 | 10.27 | 10.78 | 11.39 | 11.58 | 10.16 | 10.85 | 11.58 |

\*The increased content is partially due to the decreased reconstitution volume to 0.7 mL at T28 to add adjuvant

